# miRNA-mediated cell-to-cell communications boost DNA repair during the Radioadaptative Response

**DOI:** 10.64898/2025.12.03.692007

**Authors:** María del Carmen Domínguez-Pérez, María Jesús Fernández-Ávila, Lourdes González-Vinceiro, Laura Zannini, Héctor Peinado, Román González-Prieto, Néstor García-Rodriguez, Pablo Huertas

## Abstract

The Radioadaptive Response (RAR) is a phenomenon where a low, or priming, dose of ionizing radiation enhances cellular resistance to subsequent higher doses. We investigated whether RAR involves alterations in Homologous Recombination (HR), a high-fidelity DNA repair pathway. Using fibroblast models, we found that primed cells exhibit accelerated DNA end resection, an initial and essential HR step. This effect is mostly mediated by a bystander mechanism involving small extracellular vesicles (sEVs), as conditioned media fully replicated it. RNA profiling of sEVs identified miR-126-3p and miR-451a as key regulators of this response. Significantly, inhibiting miR-451a induced RAR in normally unresponsive cells. We further identified a miR-451a–p38–CCAR2 axis that enhances HR through suppression of CCAR2. These findings delineate a novel miRNA-mediated, sEV-driven mechanism that regulates HR during RAR, with potential therapeutic implications.

**Significance Statement:** We uncover a previously unrecognized mechanism by which human fibroblasts enhance DNA double-strand break repair through homologous recombination following a priming dose of ionizing radiation-a phenomenon known as the radioadaptive response (RAR). We demonstrate that this enhanced repair capacity is driven by small extracellular vesicle (sEV)-mediated intercellular communication, through a transient reprograming of the DNA repair capacity of neighboring cells by modulating the levels of two key microRNAs. These provide new clues on how extracellular RNA signaling governs genome maintenance, with significant implications for genome stability in healthy and pathological context. The identification of actionable modulators further strengthens the translational potential of our work.

## INTRODUCTION

Cells exposed to a source of ionizing radiation, such as those used in radiotherapy, accumulate a wide range of genomic lesions (*1*, *2*). Radiation causes various types of DNA damage, including base loss or base modifications, but primarily generates DNA breaks—most often single-strand breaks (SSBs) and less frequently, double-strand breaks (DSBs) (*2*). The vast majority of these lesions can be repaired very quickly and efficiently, using the complementary strand as a template. However, this strategy is not possible for DSBs, making them uniquely challenging and extremely toxic to repair. Indeed, a single unrepaired DSB is potentially lethal to the cell. Unrepaired DSBs are consequently the main source of cytotoxicity associated with ionizing radiation and explain the efficiency of radiotherapy in killing tumor cells (*1*, *2*). Furthermore, their repair may lead to the loss or modification of genetic information, resulting in mutations. The outcome depends on several factors, including the specific repair pathway utilized. Therefore, it is no surprising that cells have developed a complex and essential signaling network – known as the DNA Damage Response (DDR) - that is activated when DSBs, including those created by radiation, appear. The DDR affects nearly every aspect of cellular function, encompassing transcription, replication, DNA repair, and metabolic alterations, and is dependent on the post-translational modifications of thousands of target proteins (*3*). Among other key aspects, the DDR include activating cellular checkpoints and promoting the mobilization and recruitment of repair factors to DSB sites (*4*).

DSBs are DNA lesions especially difficult to repair, as both strands on the DNA molecule are affected. Consequently, there are multiple mechanisms for repairing DSBs, each with distinct requirements and outcomes (*4*). Generally, DSB repair mechanisms are classified into two broad categories based on their requirement for a homologous template. Non-homologous end joining (NHEJ) involves the direct ligation of both DNA ends, either with no processing or with minimal processing in case of “dirty” ends (*5*). NHEJ is the simplest, fastest, and energetically favorable route for DSB repair and accounts for the majority of DSB repair in mammalian cells. However, this process lacks proofreading activity, often resulting in mutations at the ligation site, ranging from minor sequence alterations to large genomic rearrangements, such as translocations, if non-adjacent DNA ends are mistakenly ligated. Thus, NHEJ is generally considered an inherently error-prone repair mechanism (*5*). In contrast, homologous recombination (HR) take advantage of the presence of homologous sequences, preferentially the sister chromatid or the homologous chromosome, and use them as templates to minimize mutagenesis of the repaired sequence (*6*). So, HR is considered mostly error-free. Within the DDR framework, a dedicated network regulates the choice between HR and NHEJ to maximize genomic stability. This regulation largely depends on the activation of the initial step in all recombination pathways: DNA end resection, a 5’ to 3’ degradation of one DNA strand at the broken end that leaves long tails of single stranded DNA (*7*). DNA resection is critical for HR and simultaneously blocks NHEJ.

A long-established but little understood phenomenon in radiation biology is the <u>R</u>adio<u>a</u>daptative <u>R</u>esponse (RARThe RAR is activated after a brief exposure to a low, non-lethal dose of radiation (the priming dose), which makes the cells resistant to a subsequent, higher, normally lethal dose (the challenging dose) for a short time (*8*, *9*). First reported in human lymphocytes in the 1980s (*10*), the RAR has since been detected in all eukaryotes, from yeast to animal models (*8*, *9*). Despite decades of study, the molecular basis of how the RAR is activated, maintained, and established remains unclear. It is known to require the activation of p53 (*8*, *9*) and p38 (*11*) and acts not only locally in the primed cells but also in neighboring, unchallenged cells via the Radiation-induced Bystander Effect (RIBE) (*12*, *13*). This cell-to-cell communication occurs both locally, through GAP junctions, but also extend farther through secreted, yet unidentified, messenger molecules (*9*, *13*). Regarding repair, it has been shown that for the triggering of the RAR an initial activation of the NHEJ, but not HR, pathway is required after the priming dose (*8*, *14*). However, the identity of the repair pathways activated in response to the subsequent, larger challenging dose is less definitively established, leading to contradictory results in the literature (*8*, *12*, *14–18*).

Considering these precedents, we studied the RAR and how it influences the repair of DSBs generated by a challenging dose. We observed that the RAR is not a universal mechanism, as not all cells can activate it. Specifically regarding repair, RAR activation leads to an acceleration of DNA end resection in response to the challenging dose. Strikingly, this response primarily depends on a bystander effect mediated by secreted messengers contained within small extracellular vesicles (sEVs). Here, we identify the differential cargo of these sEVs and demonstrate the relevance of the transient depletion of two specific miRNAs in upregulating resection after low-dose exposure. Indeed, the depletion of miR-451a is sufficient to enable the activation of the radioadaptative response in cell lines that are naturally non-responsive. Furthermore, our data suggest that DNA end resection, and consequently HR, is typically restrained due to the presence of these miRNAs, and only their reduction following a priming event triggers the activation of p38 and the inhibition of CCAR2, which is an antagonist of DNA end resection. This mechanism contributes to the transient protection of cells pre-exposed to low doses of radiation.

## RESULTS

### The RAR is cell type specific and involves an acceleration of DNA repair

To investigate whether the RAR **involves a change** in the homologous recombination pathway, we first established the experimental conditions for our study. Consistent with prior literature, immortalized fibroblasts exposed to a low dose of IR (the priming dose of 0.2 Gy) were more resistant to a larger dose (the challenging dose of 5 Gy) received 6 hours later (Figure 1A-B). However, this response was not universal, as immortalized epithelial cells did not display the same behavior (Figure 1C-D). Therefore, we chose BJhTERT cells as our model system, utilizing the 0.2 Gy priming dose and the 5 Gy challenging dose administered 6 hours later, unless otherwise specified. Some authors have speculated that the priming dose does not alter repair kinetics but instead activates an unknown protective mechanism on the DNA, leading to the creation of fewer DSBs than expected by the challenging dose (*18*). We addressed this by confirming that the enhanced survival was not due to a reduction in the amount of DNA breaks caused in primed cells when challenged. Similar levels of breaks were measured by neutral comet assay in both primed and non-primed cells following the challenging dose (Supplemental Figure 1A). Moreover, the basal number of breaks induced by the priming dose were already repaired by the time of the second irradiation (Supplemental Figure 1B).

**Figure 1.**
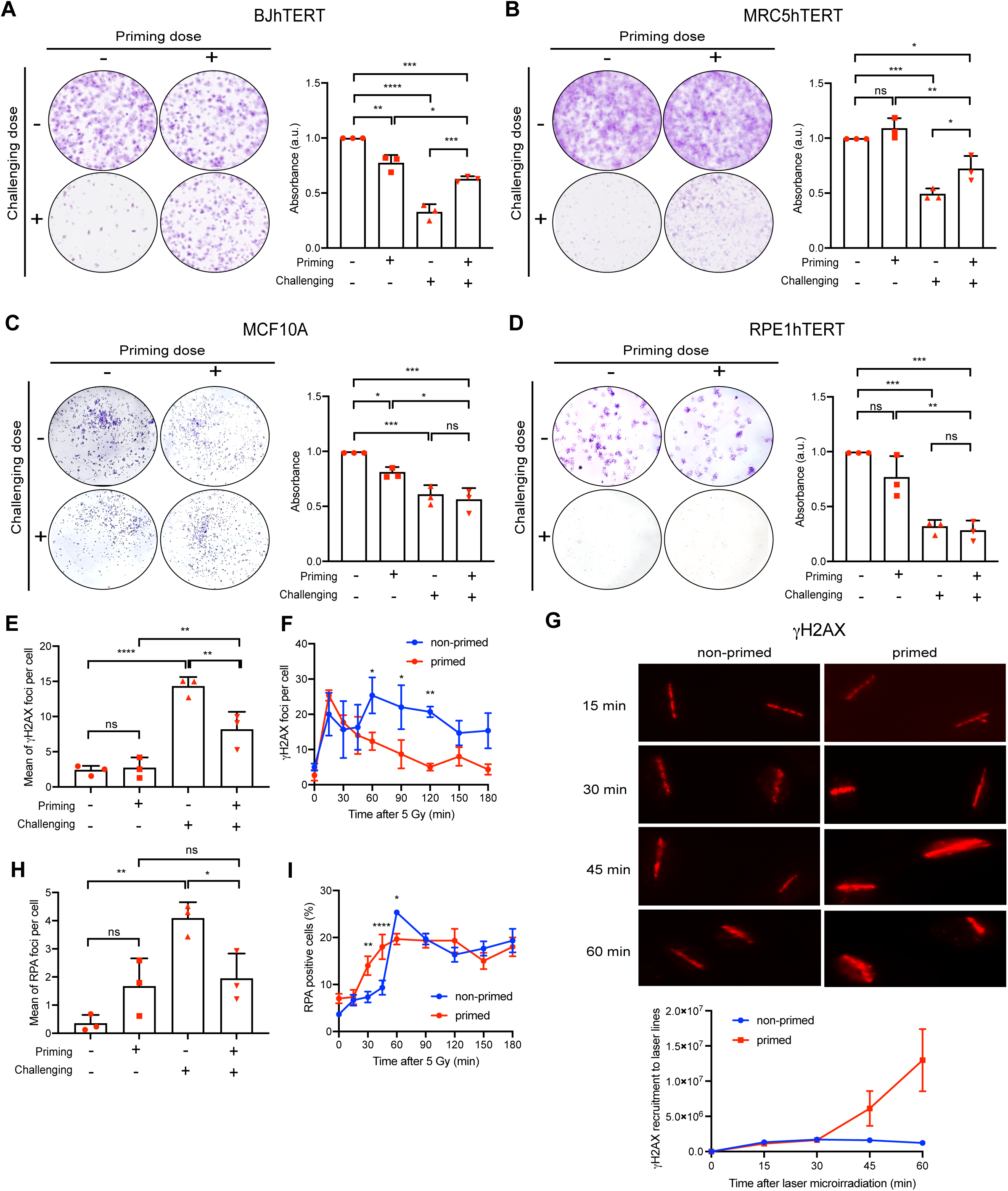
Characterization of the Radioadaptative Response in different cell lines. **A**, BJhTERT cells seeded at low density were exposed or not to a priming dose of 0.2 Gy and/or a challenging dose of 5 Gy as indicated. After forming colonies, they were stained using crystal violet (left). The dye from each well was recovered as indicated in the methods section and survival was calculated measuring the absorbance at 595 nm normalized to the cells that were not exposed to any irradiation, taken as 1. The average and standard deviation from three independent experiments are shown (right). Individual symbols in red represent individual replicas of the experiment. Statistical significance, calculated with an ANOVA test is shown as * (p<0.05), ** (p<0.01), *** (p<0.001)) or ****(p<0.0001). **B**, same as A but in MRC5hTERT cells. **C**, same as A but in MCF10A cells. **D**, same as A but in RPE1hTERT. **E**, BJhTERT cells treated as indicated in 1 were immunostained for γH2AX 1 h after the challenging dose. The number of foci per cell was scored and the mean number of foci per cell was calculated. The graph represents the average and standard deviation from three independent experiments. For each biological replica, at least 200 cells were analysed. Statistical significance was calculated as in A. **F**, BJhTERT cells exposed (primed, red line) or not (non-primed, red line) to a priming dose were challenged with 5 Gy 6h after. Cells were then collected at the indicated timepoints and immunostained for γH2AX. The mean and standard deviation from three independent experiments are plotted. For each biological replica, at least 200 cells were analysed. Statistical significance was calculated using a Student’s t-test comparing samples collected at the same timepoint. Other details as in A. **G,** BJhTERT cells, primed or not as in F, were laser microirradiated and immunostained for γH2AX at the indicated times (top). The average and standard deviation of the signal intensity from three independent experiments are plotted (bottom). For each biological replica, at least 50 cells were analysed. **H**, same as E but immunostained for RPA. **I**, same as F but immunostained for RPA.

We then monitored the repair of those breaks using histone H2AX phosphorylation (γH2AX) as a surrogate (*19*). Indeed, primed cells displayed a lower quantity of γH2AX foci per cell compared to non-primed cells 1 h after the challenging dose (Figure 1E). This reduction could reflect either faster repair kinetics or a slower signaling cascade. To resolve this, we performed kinetic analysis. Figure 1F shows that H2AX phosphorylation was similar at early timepoints, irrespective of the priming event.

However, the disappearance of the foci was significantly faster in primed cells (Figure 1F). Interestingly, although a steep reduction on the number of foci per cell was observed, the remaining ones were much brighter, suggesting an accelerated and generally stronger DDR activation (Supplementary Figure 1C). This was confirmed by using laser microirradiation as a challenging dose, which showed a much stronger accumulation of γH2AX signal at laser tracks (Figure 1G). These findings collectively support an increase in repair kinetics.

So, we wondered if during the RAR, there might be a stimulation of the most conservative but slower DSBs repair pathway, namely HR. We analyzed this by using RPA foci formation as a proxy for DNA end resection, the initiating event of HR (*7*). Similar to the γH2AX, the number of RPA foci per cell was clearly reduced 1 hour after the challenging dose (Figure 1H). Kinetic analysis (RPA foci kinetics) confirmed that this reduction was due to a speed boost rather than a slower kinetic in resection efficiency (Figure 1I). Since cell cycle is a primary regulator of DNA resection and HR, we investigated whether the priming event triggered a G2 cell cycle arrest. No changes were observed upon either the priming or challenging events 1 hour after the challenge (Supplemental Figure 1D). We also examined the NHEJ pathway using 53BP1 as a surrogate marker, but this pathway appeared unaffected during the RAR in our setup (Supplemental Figure 1E). Therefore, we conclude that during the RAR in BJhTERT fibroblasts, there is a stimulation of the HR pathways, which potentially explains the improved capability of primed cells to handle the challenging dose and thus exhibit improved survival.

### The stimulation of resection during the RAR is mediated by small Extracellular vesicles

It has been shown that some parts of the RAR rely on cell-to-cell communication either via the GAP junctions or through the cargo of small Extracellular Vesicles (sEVs), the so-called Radiation-Induced Bystander effect (RIBE) (*13*). Thus, we decided to test if this aspect of the RAR, i.e. the stimulation of resection, was controlled by either of those types of cellular communication. First, we blocked GAP junction communication using lindane (*12*) and observed that the RAR was abolished (Supplementary Figure 2A). Regarding sEVs, we hypothesized that if they were involved, exchanging the media between primed and non-primed cells right before the challenging dose (Figure 2A), might have an impact in the observed phenotypes. Indeed, swapping the media just 1h before the challenging dose not only abolished but completely reversed the pattern of RPA foci formation. Specifically, non-primed cells exposed to media from primed cells behaved identically to primed cells that retained their own media, and vice versa (Figure 2B). This effect was independent of cell cycle changes caused by the media interchange (Supplemental Figure 2B). Crucially, the media alone appeared responsible for 100% of the response, with little residual effect from whether the actual recipient cells had been primed or not (Figure 2B). This strongly suggested a secreted messenger in the media mediated the acceleration of resection.

**Figure 2.**
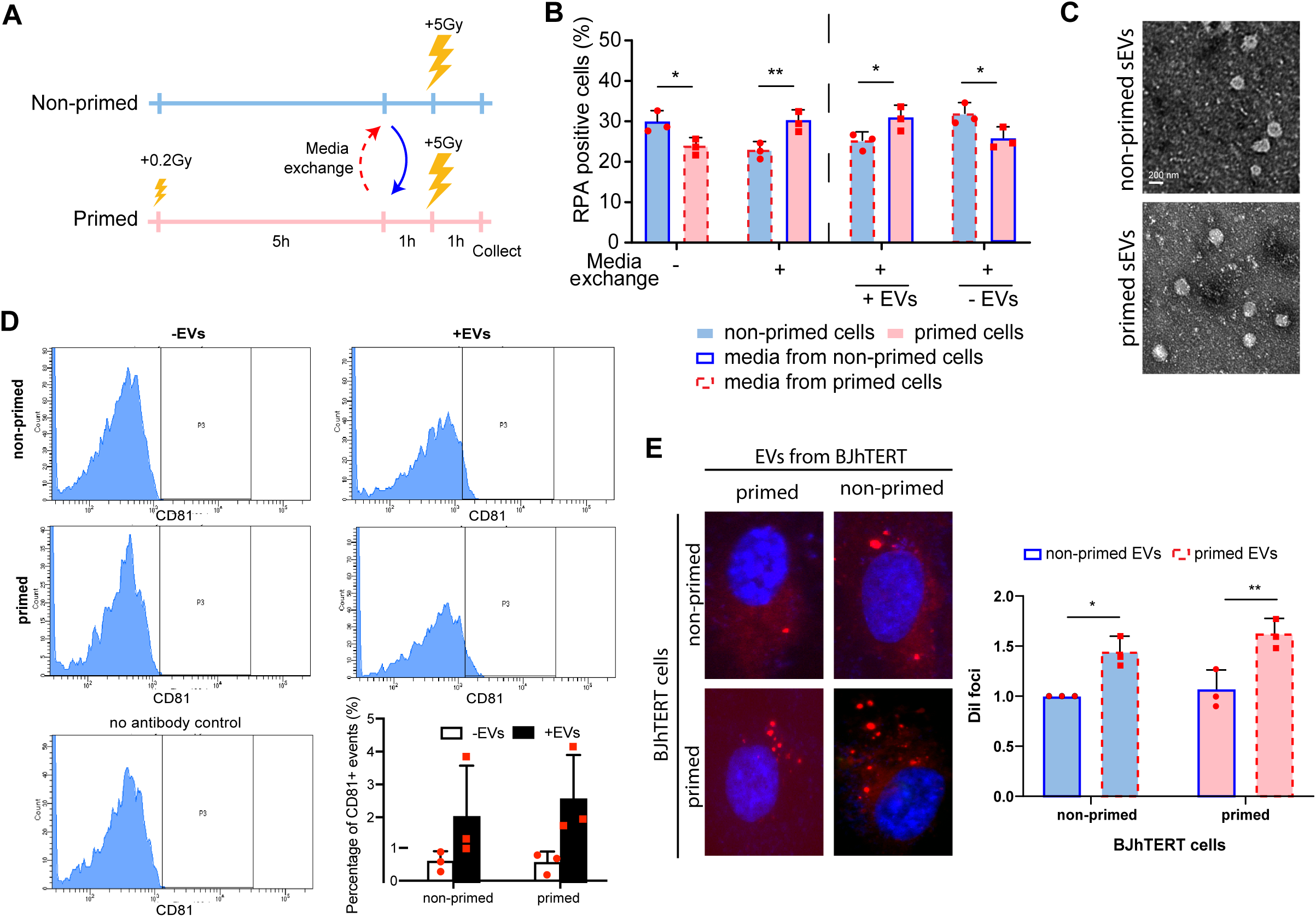
Resection boost during the RAR is mediated by sEVs. **A**, Schematic representation of the experimental setup. BJhTERT cells primed (pink) or not (cyan) were left to grow for 5h. Then, media was collected and swapped as indicated. 1h after, cells were challenged with 5 Gy and samples were collected after an additional hour for immunofluorescence. **B**, BJhTERT cells primed (pink bars) or not (cyan bars) exposed to media from either primed (dashed red border) or non-primed (solid blue border) cells were immunostained for RPA 1h after the challenging dose (left side). On the right side, the same but the collected media was either concentrated for sEVs (+EVs) or depleted of them (-EVs). Statistical significance was calculated using an ANOVA test, but only the significance of pairs of primed vs non-primed cells is shown for clarity. The average and standard deviation from three independent biological replicates are shown, and the individual values of each replica are plotted as red symbols. **C**, Representative electron microscopy images of sEVs isolated from primed and non-primed BJhTERT cells, as indicated. Scale bar, in white, represents 200 nm. **D**, Media from primed and non-primed cells (as indicated) was depleted (-EVs) or concentrated (+EVs) for sEVs was collected. Samples were stained using a CD81 antibody and the signal was measured by FACs. Representative FACs images of each category (top) and a control without antibody (bottom left), used to set the signal baseline, are shown. The average and standard deviation from three independent experiments, plus the individual replica values as red symbols, are shown on the right bottom side. **E**, BJhTERT cells treated as shown in Supplemental figure 2J were analyzed under the microscope for Dil foci. Representative images are shown on the left, and the average and standard deviation of three independent results, plus the individual values for each replica as red symbols, were plotted (right). Statistical significance was calculated as indicated in B.

To reinforce this idea, we tested an earlier media exchange. We hypothesized that exchanging media earlier would result in a mixture of the messenger present in the media and that secreted by the cells, leading to dilution of the message. We therefore expected an intermediate response in both primed cells exposed to non-primed media and non-primed cells exposed to primed media. This intermediate response was precisely what we observed (Supplemental Figure 2C). We thus concluded that while GAP junctions might play a partial role, the stronger effect occurs through a bystander effect mediated by something secreted in the media.

We then investigated whether cells unable to trigger the RAR, such as RPE1hTERT, failed due to an inability to produce the messenger, to respond to it, or both. Using BJhTERT and RPE1hTERT cells and exposing them to media from either cell type, primed or not (Supplemental Figure 2D), we found that BJhTERT cells were not affected by the addition of RPE1hTERT media. In this instance, the reduction in RPA foci 1 hour after the challenging dose was still observed even when primed BJhTERT cells were exposed to media from non-primed RPE1hTERT cells, likely due to the accumulation of the messenger secreted by the BJhTERT cells themselves in the final hour before the challenge. Conversely, there was no reduction in RPA foci number in RPE1hTERT cells regardless of the media used. We therefore concluded that RPE1hTERT cells can neither emit an intelligible messenger nor respond to it to trigger this effect.

Next, we tested if the messenger in the media was contained within the cargo of sEVs. To do so, we purify these vesicles from the conditioned media as described in the methods section. Strikingly, adding the sEV fraction alone was sufficient to replicate the response observed when swapping the whole media (Figure 2B, Supplementary Figure 2C). Conversely, when the media was depleted of sEVs, the cells retained the behavior dictated by their own irradiation history (Figure 2B, Supplementary Figure 2C). This strongly suggests sEVs are responsible for this specific response. Indeed, sEVs were capable of completely overriding the response when present, meaning that the outcome depends solely on the nature (primed or not) of the cells that produced the sEVs, high jacking the signal produced by the actual challenged cells. When sEVs were depleted, the pre-irradiation history of the cells prevailed, showing the RAR if the cells themselves were primed.

### sEVs from primed cells can be internalized faster than those form non-primed cells

We further characterized the sEVs secreted from primed versus non-primed BJhTERT cells. Electron microscopy was used to examine the physical characteristics of the sEVs (Figure 2C). Analysis showed no changes in diameter or area, and only minor changes in roundness and perimeter (Supplemental Figure 2E-H). We analyzed typical sEV markers, CD81 and CD9, using FACs with specific antibodies, noting that the positivity for these markers varies based on sEV composition (20). We established a baseline using sEV-depleted media and then analyzed the purified sEVs (Figure 2D, Supplemental Figure 2I). We observed vesicles decorated with CD81, but not CD9, in the sEV-retaining fraction, which were absent in the sEV-depleted samples (Figure 2D, Supplemental Figure 2I). However, there was no major difference in the quantity of CD81-positive sEVs between primed and non-primed samples (Figure 2D).

Next, we studied their internalization using the Dil labelling (see Method section in the supplementary information). Briefly, primed and non-primed cells were incubated with the Dil label for 1h, then the media was collected and sEVs purified and added to a fresh set of primed or non-primed cells that were Dil free (Supplemental Figure 2J). sEVs from primed cells were found to be more avidly internalized than sEVs from non-primed cells, regardless of the status of the recipient cells (Figure 2E). This suggests that a specific subpopulation of sEVs in the primed media exhibits higher uptake. To determine whether the unresponsiveness of RPE1hTERT cells to these sEVs is due to their inability to detect and internalize this sEV subpopulation, we repeated the experiment using Dil-labeled sEVs exchanged between BJhTERT and RPE1hTERT cells (Supplemental Figure 2K-L). In both scenarios, RPE1hTERT cells failed to increase sEV uptake, suggesting not only that they are "deaf" to the RAR-related sEVs from BJhTERT, but also that they cannot produce them .

### RAR-mediated resection stimulation is mediated by miRNA

We hypothesized that changes in the composition of the cargo within the primed sEVs might account for their impact on the RAR-mediated boost of DNA repair. Changes in protein, RNA, and DNA content following DNA damage exposure have been reported previously (*21*). We compared the composition of primed versus non-primed sEVs using mass spectrometry for protein cargo (Supplementary Dataset 1) and total RNA sequencing for RNA cargo (Supplementary Table 1, Figure 3A). Reproducible changes were confirmed in both cargo types. Strikingly, only a few RNAs were differentially present in primed versus non-primed sEVs (Figure 3A, Supplementary Table 1), and only a subset of these were linked to the response to ionizing radiation (Figure 3B) . Among them, two miRNAs stood out due to loose associations with the DNA damage response: miR-126-3p (*22–24*) and miR-451a (*25*) (Figure 3A, Supplementary Table 1).

**Figure 3.**
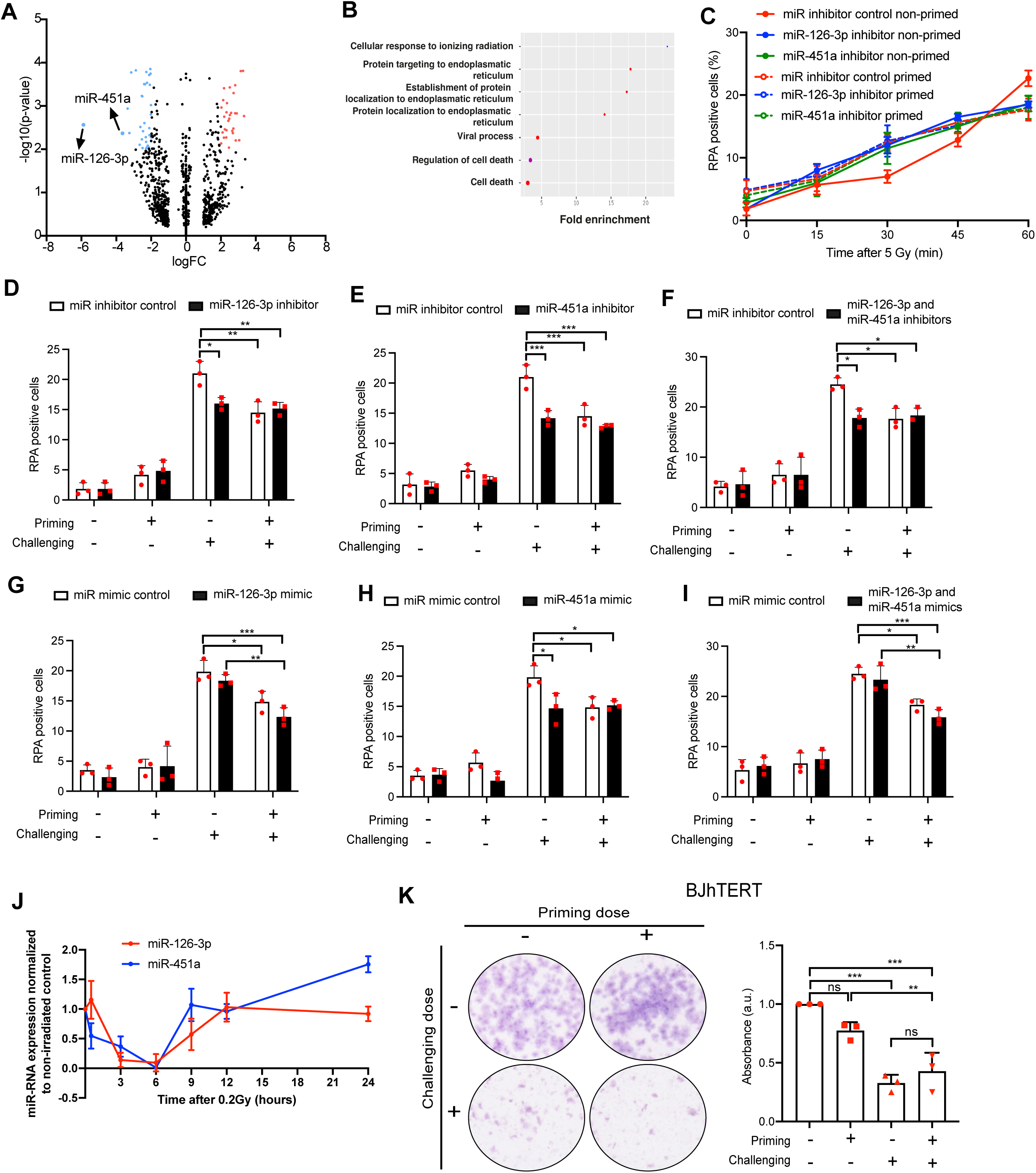
miR-451a and miR-126-3p downregulation upon a priming event mediates the RAR. **A**, Volcano plot representing the fold-change abundance of RNAs present in sEVs of primed versus non-primed BJhTERT. Red dots represent those enriched over the statistical significance in primed sEVs. In blue, those depleted on those circumstances. miR-126-3p and miR-451a are highlighted. **B**, GO analysis of the result showed in A and Supplementary Table 1. **C**, BJhTERT cells exposed or not to the priming dose and transfected with different miRNA inhibitors, as indicated, were challenged with 5 Gy. Samples were taken at the indicated timepoints and immunostained for RPA. The average and standard deviation from three independent replicas are plotted. **D**, BJhTERT transfected with a miRNA inhibitor against miR-126-3p (black bars) or a control inhibitor (white bars), were exposed or not to the priming and challenging dose as indicated. 1 h after the challenging dose samples were taken an immunostained for RPA. The average and standard deviation of three independent biological replicates is shown, and the individual values of each replica are plotted as red symbols. Statistical significance was calculated using an two way ANOVA test, but only the significance of selected meaningful pairwise comparisons are shown for clarity. **E**, Same as D but using an inhibitor against miR-451a. **F**, Same as D but combining the inhibitors against miR-126-3p and miR-451a. **G**, Same as D, but using miRNA mimics instead of inhibitors. **H**, Same as G but using the miR-451a mimic. **I**, Same as G but combining miR-126-3p and miR-451a mimics. **J**, Expression of miR-126-3p (red) and miR-451a (blue) in BJhTERT was calculated by qRT-PCR in samples at the indicated times after the priming dose. The average and standard deviation from three independent experiments are shown. **K**, Same as figure 1A, but cells were challenged with 5 Gy 12 h after the priming dose.

miRNA contained in sEVs have been described as being involved in DNA repair (*21*). Surprisingly, we observed a reduced amount of both miRNAs in primed sEVs when compared with non-primed. We tested their involvement in the RAR-mediated resection boost by first reducing their presence using miRNA inhibitors. Our expectation, based on their reduction during priming, was that their artificial reduction would trigger the RAR response even in non-primed cells. Indeed, inhibition of either miRNA independently resulted in an acceleration of resection that perfectly mimicked the RAR observed in control cells exposed to a priming dose (Figure 3C-E). This phenotype involved an increase in RPA foci at earlier time points, followed by a decrease 1 hour after the challenging dose. Notably, when either miRNA was inhibited, no further effect was observed upon exposing the cells to a priming dose, strongly supporting the idea that their reduction is essential for resection stimulation. Combining both inhibitors did not further affect resection and yielded the same phenotype in non-primed cells as priming with 0.2 Gy (Figure 3F) . Overexpression of either or both miRNAs was not sufficient to abolish the RAR response (Figure 3G-I), indicating that while the reduction of either miRNA is sufficient to trigger the response, additional elements are involved during the RAR.

Since the RAR is a transient protection of the cell against IR, we correlated this temporality with miRNA expression. Kinetic analysis showed that both miRNAs rapidly declined after the priming event, reaching a nadir 6 hours later (Figure 3J). This was followed by recovery over the next 6 hours, with expression completely recovered 12 hours after the priming event (Figure 3J, 281). In agreement with our model, when primed BJhTERT cells were challenged 12 hours, rather than 6 hours, after the priming dose, the protective effect was lost (Figure 3K).

### miR-451a inhibition can activate the RAR in RPE1hTERT cells

We wondered if natural differences in expression of miR-126-3p and/or miR-451a might explain why some cells fail to activate the RAR. So, we first compared the levels of both miRNA by RT-qPCR in RPE1hTERT vs BJhTERT cells. Interestingly, both miRNAs exhibited distinct expression profiles between the cell lines. RPE1hTERT cells naturally displayed very low levels of miR-126-3p but strongly expressed miR-451a (Figure 4A-B). Furthermore, priming RPE1hTERT had divergent consequences for the two miRNAs. While the basally low miR-126-3p was increased upon priming, miR-451a levels decreased in a manner similar to that observed in BJhTERT cells (Figure 4C-D, Supplemental Figure 3A-B).

**Figure 4.**
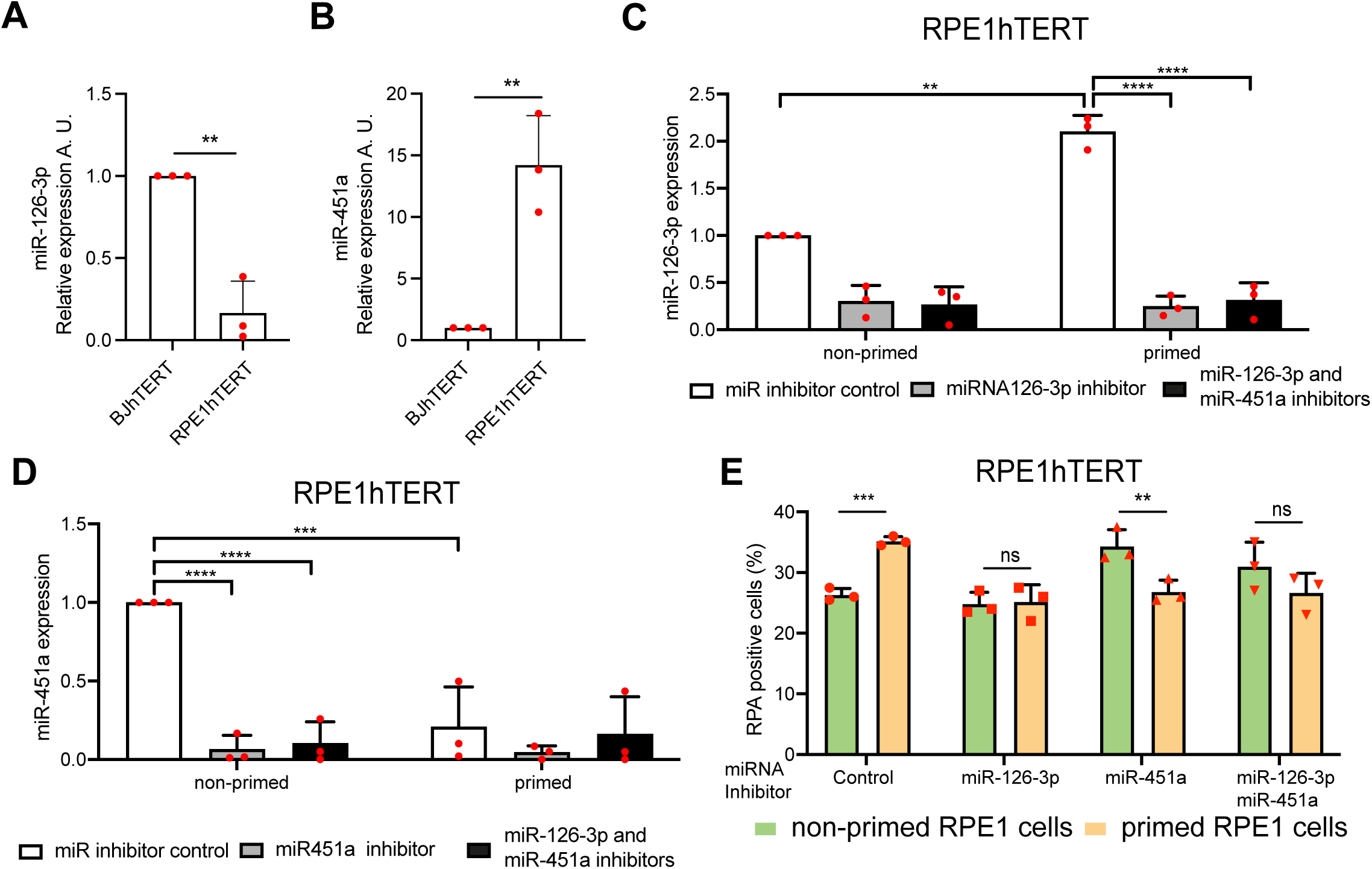
miR-451a depletion activates the RAR in RPE1hTERT cells. **A**, Expression of miR-126-3p in BJhTERT cells or RPE1hTERT cells, as indicated, was calculated by qRT-PCR. Expression was normalized to the BJhTERT results, taken as 1. The average and standard deviation from three independent experiments are show, and individual results are depicted as red symbols. Statistical significance was calculated using a Student’s t-test. **B**, Same as A but for miR-451a expression. **C**, Same as A, but in cells primed or not as indicated and transfected with miRNA inhibitor against miR-126-3p (grey bars), both miR-126-3p and miR-451a (black bars) or a control (white bars). In this case, statistical significance was calculated using a two-way ANOVA. **D**, Same as C but for cells transfected with an miR-451a inhibitor (grey bars). **E**, RPE1hTERT cells transfected with the indicated miRNA inhibitors and primed (yellow bars) or not (green bars) were collected 1h after the challenging dose and immunostained for RPA. Other details as figure 2B.

Thus, we hypothesized that this profile explains why RPE1hTERT cells fail to trigger the response. In this scenario, miR-126-3p would have little effect due to its upregulation following the priming dos . miR-451a might be relevant, but its high basal levels could keep it above a necessary threshold even after priming. If this were the case, miR-126-3p inhibition should have minimal effect in RPE1hTERT cells, whereas artificial inhibition of miR-451a could cause the levels to drop below the hypothetical threshold following low-level radiation exposure. Indeed, miR-451a, but not miR-126-3p, inhibition restored the RAR in RPE1hTERT cells (Figure 4E). Double inhibition of both miRNAs also restored the RAR, likely due only to miR-451a effect. Thus, depleting a single miRNA was sufficient to activate the RAR in cells that do not naturally exhibit it. However, and consistent with the lack of ability of RPE1hTERT cells to increase the uptake of sEVs, even when cells were treated with the miR-451a inhibitor, they were unaffected by conditioned media from either inhibitor-transfected RPE1hTERT or BJhTERT cells (Supplementary figure 4A-B).

### Resection boost during the RAR is mediated by p38-CCAR2

We sought to identify the internal factors mediating the RAR-induced boost of DNA end resection. Mass spectrometry analysis of sEV proteins revealed only three proteins differentially loaded when primed sEVs were compared with non-primed sEVs (Supplementary dataset 1). Among these was MAPK1/ERK2, a regulator of the p38 stress-induced pathway. Furthermore, miR-451a’s role in DNA repair has been shown to depend on this pathway (*25*). Furthermore, p38 has been shown to be implicated in other aspects of the RAR (*11*). We therefore hypothesized that sEV-mediated regulation of MAPK signaling might regulate resection. To test this idea, we first studied p38 activation by the priming dose or the challenging dose (Figure 5A). Indeed, only the priming dose stimulated p38 activity 1 hour after irradiation (Figure 5A; Supplementary figure 5A) . Using the p38 general inhibitor losmapimod, we confirmed that a single priming event maintained p38 activity for hours, even at the time of the challenging dose, whereas the higher challenging dose did not activate this response (Supplementary Figure 5B). Moreover, no further p38 activation was observed when the priming and challenging doses were combined (Figure 5A). Then, we found that p38 inhibition with losmapimod deactivated the resection boost associated with the RAR (Figure 5B). This drug inhibits two out of the four isoforms of p38, namely p38α and p38β. To determine which isoforms were required for this process, we depleted either one using siRNA. As shown in figure 5C, only p38α downregulation completely abolished the RAR effect on DNA end resection, while p38β depletion rendered an intermediate phenotype. No changes in cell cycle could explain this effect (Supplementary Figure 5C). Strikingly, p38 inhibition not only abolished the consequences of the priming dose on RPA foci but was epistatic over the effect of conditioned media (Figure 5D), suggesting that p38 activation, both in the emitting and recipient cells, is essential for the bystander effect.

**Figure 5.**
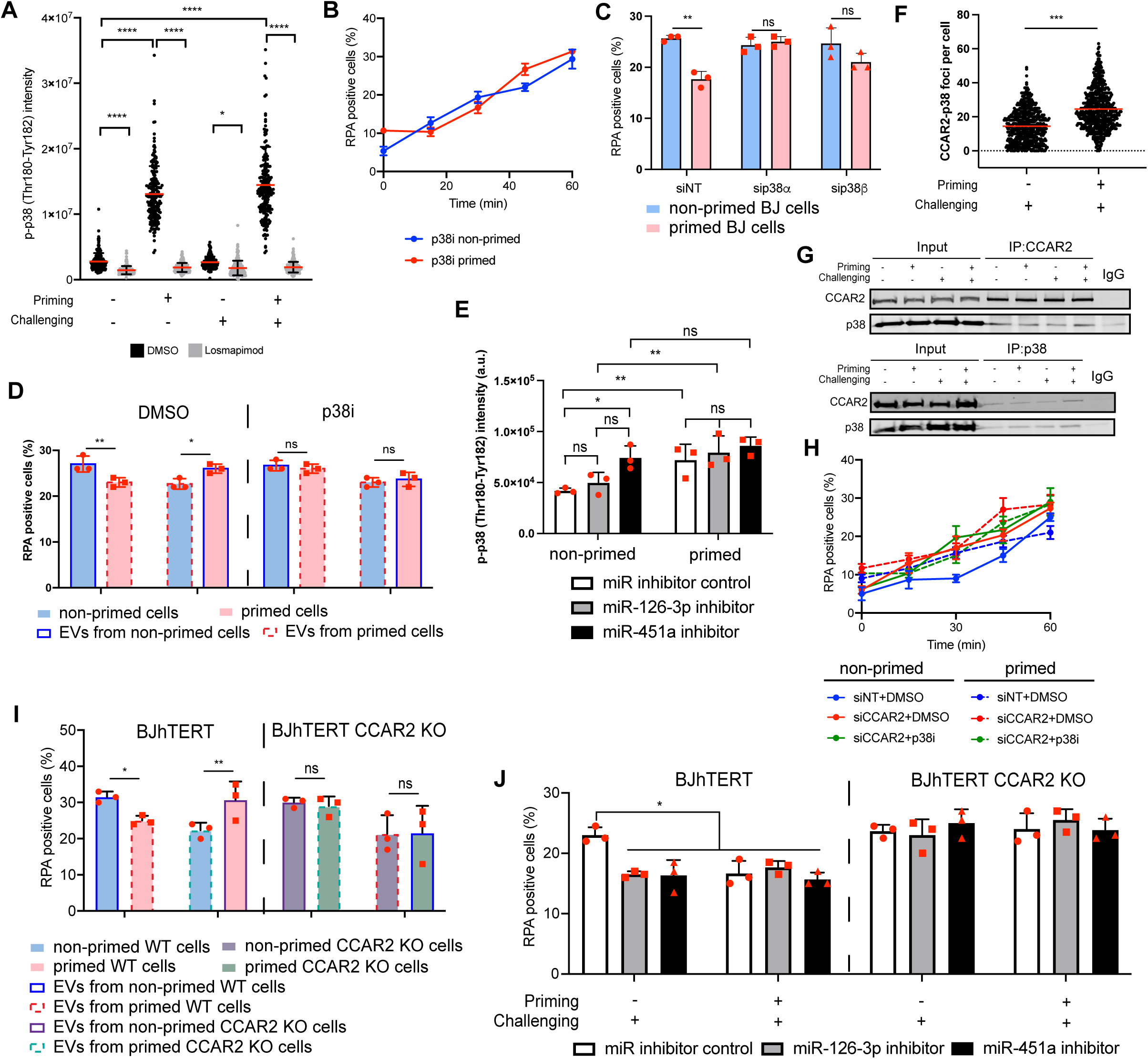
Resection boost during the RAR is mediated by p38 activation and CCAR2 inhibition. **A**, BJhTERT cells were treated with the p38 inhibitor losmapimod (grey) or DMSO as control (black) and exposed to a priming and/or challenging dose as indicated. Then, cells were immunostained using an antibody against phosphorylated p38. The signal intensity per cell was scored automatically and plotted. Statistical significance was calculated using a two-way ANOVA test. Only pairwise comparison between similar samples treated with DMSO or losmapimod are shown for clarity. **B**, BJhTERT cells treated with losmapimod and exposed (primed, red) or not (non-primed, blue) to the priming dose were challenged with 5 Gy. Samples were taken at the indicated timepoints and immunostained for RPA. The average and standard deviation from three independent replicas are plotted. **C**, BJhTERT cells transfected with the indicated siRNAs and primed (pink bars) or not (cyan bars) were collected 1h after the challenging dose and immunostained for RPA. Statistical significance was calculated using a two-way ANOVA test, but only the significance of pairs of primed vs non-primed cells is shown for clarity. The average and standard deviation from three independent biological replicates are shown, and the individual values of each replica are plotted as red symbols. **D**, BJhTERT cells primed (pink bars) or not (cyan bars) exposed to media from either primed (red dashed border) or non-primed (blue solid border) cells were immunostained for RPA 1h after the challenging dose. Cells were either exposed to DMSO (left side) or the p38 inhibitor losmapimod (right side). Other details as in C. E, same as A, but in cells transfected with the indicated miRNA inhibitors and after exposure or not to a priming dose. **F**, Cells primed or not were exposed to the challenging dose 6h afterwards and immunostained using a PLA method as described in the methodology section using p38 and CCAR2 antibodies. The number of CCAR2-p38 PLA foci per cell from three different experiments was automatically quantified and plotted. Statistical significance was calculated using a Student t-test. **G**, Protein samples from BJhTERT cells, exposed or not to the priming and challenging doses, were used for immunoprecipitation using an anti-CCAR2 (top) or p38 (bottom) antibody. Input (left) and immunoprecipitated (right) samples were resolved in SDS-PAGE and immunoblotted using both antibodies. A representative experiment is shown. **H**, Same as B but for cells primed (dashed lines) or not (solid lines) exposed to DMSO or losmapimod and/or transfected with an siRNA control or against CCAR2, as indicated. The average and standard deviation from three independent experiments are shown. **I**, Wildtype (left) or CCAR2 KO (right) BJhTERT cells were primed or not as indicated and cultured with media from primed or not primed wild type or KO cells as labelled. Other details as in D. **J**, Wildtype (left) or CCAR2 KO (right) BJhTERT cells transfected with the indicated miRNA inhibitors were primed or not as indicated. 1h after the challenging dose, samples were immunostained for RPA. Other details as in D.

To test whether p38 activation during the RAR was caused by the depletion of miR-126-3p and/or miR-451a, we quantified kinase phosphorylation upon miRNA inhibition. Strikingly, activation of p38 was observed upon inhibition of miR-451a regardless of exposure to the priming dose (Figure 5E), mimicking the effect of the priming dose alone (Figure 5A). Little effect was observed upon miR-126-3p inhibition, as p38 activation in that case relied solely on the priming event. We thus concluded that p38 activation during RAR relies on the depletion of miR-451a.

We then investigated which protein might be targeted by p38 to accelerate resection during the RAR. It has been shown that the anti-resection protein CCAR2 is dephosphorylated by WIP1, which eliminates phosphate groups dependent on this stress kinase (*27*). Thus, we hypothesized that during the RAR, p38 activation might inhibit CCAR2, thereby boosting resection. First, we demonstrated that CCAR2 was required for the increased survival associated with the RAR (Supplementary Figure 5D). We then tested if p38 and CCAR2 physically interact and if this interaction was modulated by the priming event. Proximity Ligation Assay (PLA) showed that they were in close proximity in challenged cells, and even more so if the cells were initially primed (Figure 5F). Furthermore, reciprocal co-immunoprecipitations showed a basal interaction between CCAR2 and p38 that was slightly induced specifically in cells that were both primed and challenged (Figure 5G). To functionally link these proteins, we tested resection kinetics in cells depleted or not for CCAR2 and inhibited or not for p38 with losmapimod (Figure 5H) . In agreement with our hypothesis, CCAR2 depletion not only increased resection, as previously described, but closely mimicked the boost associated with the RAR . Moreover, there was no further boosting in CCAR2-depleted cells when the RAR was triggered by a priming event, and p38 inhibition had no effect in this background (Figure 5H).

Finally, we explored whether CCAR2 was indeed the central component upon which sEV regulation converges to control resection during the RAR. In this scenario, cells lacking CCAR2 should still be able to produce sEVs but should be unable to react to them. This is what we observed (Figure 5I). To functionally connect the downregulation of miR-126-3p and/or miR-451a with CCAR2, we checked the effect of miRNA inhibitors in wildtype and CCAR2 KO BJhTERT cells. Consistent with the model, inhibition of either miRNA caused a decrease in RPA foci formation in non-primed cells (mimicking the RAR), but this effect was completely lost when CCAR2 was absent (Figure 5J). We conclude that the signal contained on the sEVs produced during the RAR, specifically the downregulation of miR-126-3p and miR-451a, requires CCAR2 in the recipient cells to exert its boosting effect on DNA end resection. Our data also suggest that miR-451a acts through p38 activation, whereas miR-126-3p utilizes an alternative means that, nonetheless, also impacts CCAR2.

## DISCUSSION

The RAR is a highly complex and poorly understood stress response that is critical in defining how cells react to repeated radiation exposure. In the context of radiotherapy, this is highly relevant, as it impacts both how cancer cells respond to treatment and how healthy neighboring cells are affected. This is particularly important given the bystander effect, where cells communicate with each other. Understanding how this protective mechanism is activated and shared between cells is crucial in radiotherapy, both for maximizing the toxic effect on cancer cells and minimizing negative effects on surrounding healthy tissue. To complicate the issue further, the RAR is not a universal phenomenon, and cell types display varied responses. Understanding the nuances of different cell types will be critical in future studies. Interestingly, this variability might stem from the expression levels of only a few factors, given that we were able to reactivate the RAR in RPE1hTERT cells simply by depleting a single miRNA. Moreover, it is known that genetic components modulate the RAR differently among individuals based on their genetic makeup, as lymphocytes from different donors behave variably (*28*) and the response differs more between fraternal twins than between identical twins (*29*). These changes can be dynamic and responsive to the environment, with specific changes observed among individuals living in high natural radiation areas (*30*). Identifying the critical factors involved in the RAR will aid in understanding cell type, genetic, and environmental differences, potentially serving as biomarkers to determine if a cell can enact the RAR.

The molecular components of the RAR are still poorly understood. Additionally, there is a controversy regarding the contribution of different DSB repair pathways (*8*, *12*, *14–18*). Interestingly, our own results highlight the importance of detailed studies. Indeed, by assessing repair proficiency at a single time point (e.g., 1 hour after the challenging dose), resection might misleadingly appear downregulated. However, as revealed by kinetic analysis, this actually reflects an acceleration in DNA end processing, rather than a decrease. This necessity of checking the repair process at appropriate times may partially explain the seemingly contradictory results in the existing literature .(*8*, *12*, *14–18*).

A key breakthrough of our study is that the RAR, concerning its reliance on DNA end resection and homologous recombination, does not rely on activation of the process but rather on the transient release from a constitutive inhibition. This implies that cells, at least fibroblasts, appear to continuously suppress their own repair capacity by expressing specific miRNAs that constitutively activate CCAR2, a repressor of resection. A similar dampening of repair via CCAR2 has been previously reported (*31*, *32*). One initial, counterintuitive explanation for this model is that it likely facilitates a faster response, bypassing the need to express and accumulate resection and recombination proteins. But why do cells not maintain their repair capacity constantly at its maximum? This suggests that maintaining a heightened state of repair may be detrimental to cell fitness and survival. This could be due to the higher energetic cost of recombination compared to NHEJ. Thus, when breaks are unlikely, promoting end-joining might be more cost-effective without severely compromising genomic stability. However, in stressful situations, such as following a priming irradiation, HR may be required to guarantee genomic integrity. A non-mutually exclusive explanation is that increased resection can be toxic, as resection must be strictly limited to avoid processing stalled replication forks (*33*). Consistent with this idea, long-term depletion of either miR-126-3p or miR-451a leads to cell lethality (not shown).

Strikingly, the most relevant contributor of RAR in human fibroblast is the bystander effect through sEVs. Although cells can support the RAR independently, the presence of these sEVs is sufficient to dominate the internal response. Consequently, cells behave largely as responders dictated by the status (primed or not) of the cells that produced the sEVs in the media. In this scenario, cells globally coordinate their response via this bystander effect, relying more on this collective signal than on their own, self-produced, marks. In our model (Figure 6), fibroblasts continuously produce sEVs containing, among other factors, miR-126-3p and miR-451a. Self-produced vesicles (Figure 6A, blue circles) are rapidly exported, creating an environment rich in sEVs loaded with these miRNAs (red circles). These sEVs are internalized, maintaining a high cellular presence of the miRNAs 8Figure 6A). This homeostasis ensures that cellular miRNA levels primarily reflect the cargo of the sEVs rather than self-produced RNAs. Consequently, p38 is inactive, and CCAR2 constitutively restrains CtIP, dampening resection via both miR-451a-p38-mediated regulation and an alternative miR-126-3p pathway (Figure 6B). Upon exposure to a priming dose, irradiated cells (Figure 6C, yellow cells) modify the composition of the sEVs they produce, affecting both RNA and protein content (white circles). Among other changes, the amount of those two miRNAs is reduced in the vesicles. Increased uptake of the primed sEVs (white circles) result on a progressive, temporal depletion of miR-126-3p and miR451a within the receiving cells through dilution, causing all cells to behave as if they were primed (Figure 6D-E). This dilution activates p38 via miR-451a depletion. This, combined with a p38-independent, miR-126-3p-dependent mechanism, leads to CCAR2 inhibition (Figure 6F). The subsequent release of CCAR2 stimulates CtIP and DNA end resection (*31*, *34*), thereby boosting the cells’ repair capacity. Later (Figure 6G), cells re-activate the production of both miRNAs, which are loaded into vesicles (blue circles) and exported as sEVs (red circles). This process slowly restores the initial homeostasis, where sEVs loaded with miR-126-3p and miR-451 maintain high internal miRNA levels, switch off p38, and dampen resection through CCAR2 (Figure 6A-B).

**Figure 6.**
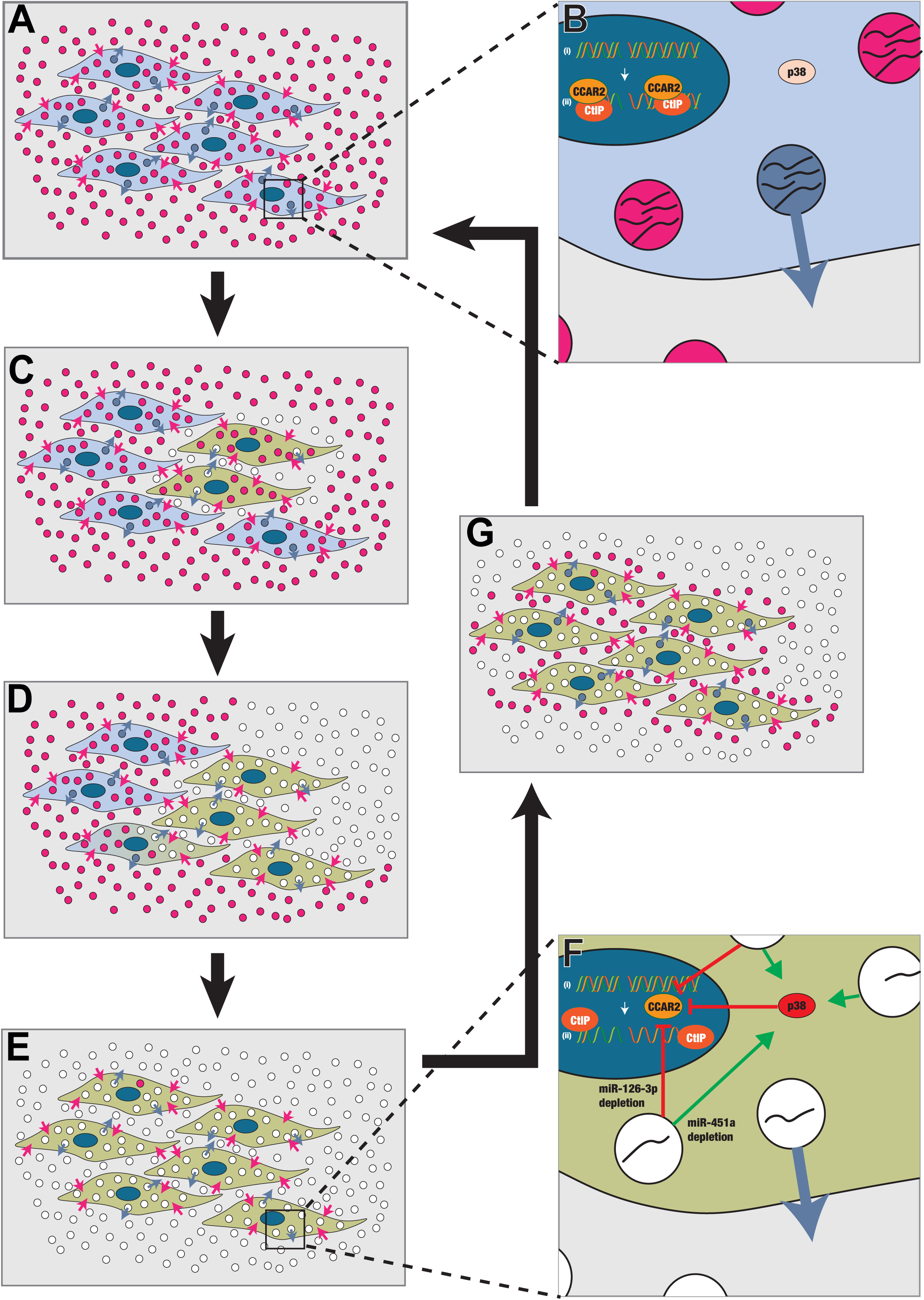
Model of the sEV-mediated effect on DNA resection during the RAR. **A**, Fibroblasts are continuously producing sEVs. Self-produced vesicles (blue circles) are quickly exported to create an environment full of sEVs (red circles), that are internalized and establish a homeostasis in which the cellular levels of the miRNAs depend mostly on the cargo of the sEVs and not the self-produced RNAs. **B**, As a response, p38 is not activated and CCAR2 is constitutively harnessing CtIP thus dampening resection. **C**, Upon the exposure to a priming dose, cells that have been irradiated (yellow cells) change the composition of the sEVs they produce, both at the RNA and protein levels (white circles). Among other changes, the amount of miR-126-3p and miR-451a is reduced in the vesicles. **D-E**, Increased uptake of the primed sEVs (white circles) result on a progressive temporal depletion of the miRNAs within the receiving cells by dilution until all cells behave as if they were primed. **F**, This dilution activates p38 through depletion of miR-451a. That, together with a miR-126-3p dependent mechanisms, inhibit CCAR2 and stimulates CtIP and DNA end resection. **G**, Later on, cells re-activate the production of both miRNAs that are loaded in vesicles (blue circles) and exported as sEVs (red circles). **A-B**, Slowly, this restores the initial homeostasis in which sEVs loaded with mIR-126-3p and miR-451 keep the internal levels of those miRNAs high, switch off p38 and dampen resection through CCAR2.

Beyond the RAR, similar sEV-mediated regulations of DNA repair might be active in other contexts. Extracellular vesicles have previously been proposed to trigger self-protective mechanisms against DNA damage in parental and recipient cells, although conflicting reports exist in the literature (*21*).

The secretion of sEVs is a crucial component of the cellular microenvironment in both physiological and pathological settings (*35*). For instance, during inflammation, cells release sEVs containing miRNAs, cytokines, and lipids, which propagate the inflammatory signal to neighboring or distant cells, thereby altering the surrounding microenvironment (*35*). Within this context, miRNAs act as key regulators of inflammation by controlling the expression of genes involved in immune signaling, cytokine production, and cell migration. Strikingly, inflammation and DSB repair are intricately linked, as inflammatory responses can influence the efficiency and accuracy of DNA repair, and DNA damage can trigger the release of inflammatory cytokines (*36*, *37*). Inflammatory signals can modulate the activity of key proteins involved in DSB repair. Moreover, chronic inflammation may lead to sustained DNA damage and inefficient DSB repair, promoting genomic instability and increasing mutation risk, potentially contributing to tumorigenesis or inflammatory-related diseases. Additionally, the persistence of unresolved DNA damage under inflammatory conditions can exacerbate the inflammatory response itself, creating a vicious cycle of DNA damage and inflammation that contributes to disease progression. Our findings may clarify how these processes are linked and why the interplay between the cellular microenvironment, inflammation, and DNA repair mechanisms is crucial for maintaining cellular health and preventing chronic diseases.

This coordinated response of cells through sEVs opens the possibility to influence the tissue reaction during irradiation. However, the nuances on genetic and environmental differences in the response need to be considered, such as the fact that RPE1hTERT cells downregulated for miR-451a, that reactivate the RAR, do so in a way that is independent of the conditioned media. This might be explained by the intrinsically high levels of self-produced miR-451a, making them less susceptible to the miRNA composition of sEVs. Furthermore, this dependence on miR-451a, but not miR-126-3p, suggests RPE1hTERT relies on the p38 branch of the regulatory mechanism. Again, this can simply be explained by the extremely low abundance of this miRNA even in non-primed conditions. Thus, further studies might provide compelling evidence on the universality of this effect. In any case, by identifying specific factors, primarily miR-451a and p38, we define biomarkers that could help predict RAR responsiveness. Moreover, the fact that depletion of any of these factors alone is sufficient to activate the RAR provides a molecular tool to induce it . In this scenario, we propose that delivering specific miRNA inhibitors to target cells could induce a protective mechanism against radiation, offering an interesting strategy to minimize unwanted toxic side effects of radiotherapy in healthy tissue . Our data may also be extrapolated to other DNA damage-induced therapies; for example, miR-126-3p expression has been shown to induce cisplatin resistance in triple-negative breast cancer (*24*).

Taken together, our results establish a mechanistic framework for understanding how the RAR enhances DNA repair by stimulating HR. We propose that priming irradiation alters sEV-mediated intercellular communication, leading to a boost in DNA repair through several independent pathways. We defined one pathway mediated by the depletion of miR-126-3p and another by the downregulation of miR-451a and subsequent p38 signaling activation. Both pathways function by releasing the CCAR2-mediated inhibition of DNA end resection, ultimately accelerating HR. These findings not only advance our understanding of RAR but also have potential implications for improving radiotherapy strategies by exploiting sEV-mediated signaling to enhance DNA repair in normal tissues while selectively sensitizing tumor cells to IR.

## Acknowledgements.

Funding: This work was funded by the R+D+I grant PID2022-136791NB-I00 funded by MICIU/AEI/10.13039/501100011033/ FEDER/UE. MdCD-P is funded with a FPI fellowship number PRE2020-091834. NG-R is funded by the EMERGIA 2021 program from the Junta de Andalucía (EMC21_00057). CABIMER is supported by the regional government of Andalucía (Junta de Andalucía). We also want to thank the Microscopy and Genomic Unit of CABIMER for their support in the experiments and Iván V. Rosado for critical reading of the manuscript.

## Author contribution

McD-P performed all the experiments with the help of MJF- A. LG-V and RG-P performed the mass spectrometry analysis of sEVs. LZ helped with the analysis of CCAR2 KO cells. HP provided help with the sEVs analysis. NG-R supervised CdD-P. PH formulated the initial hypothesis, supervised the work and analyzed the data. PH also wrote the initial manuscript.

The authors declare no conflict of interest.

## DATA AVAILABILITY

All relevant data are included in the manuscript. Raw data will be provided upon request. RNA-seq data are deposited at GEO under the accession number GSE293977 and the mass spectrometry proteomics data have been deposited to the ProteomeXchange Consortium via the PRIDE (*38*) partner repository with the dataset identifier PXD063287.

## Material and Methods

### Cell lines and growth conditions

BJhTERT, BJhTERT CCAR2-KO and RPE1hTERT cells were grown in high-glucose Dulbecco’s Modief Eagle Medium (DMEM) medium with L-glutamine (Sigma-Aldrich) supplemented with 10% fetal bovine serum (Sigma-Aldrich), 100 units/ml penicillin, and 100 μg/ml streptomycin (Sigma-Aldrich).

MRC5hTERT cells were grown in low-glucose Dulbecco’s Modief Eagle Medium (DMEM) medium without L-glutamine (Sigma-Aldrich) supplemented with L-Glutamine (Gibco), 10% fetal bovine serum (Sigma-Aldrich), 100 units/ml penicillin, and 100 μg/ml streptomycin (Sigma-Aldrich).

MCF10A cells were grown in DMEM/F-12 medium with L-glutamine (Sigma-Aldrich) supplemented with 5% donor horse serum (Sigma-Aldrich), 20 ng/ml epidermal growth factor (EGF) (Sigma-Aldrich), 10 µg/ml insulin (Sigma-Aldrich), 0.5 µg/ml hydrocortisone (Sigma-Aldrich), 100 ng/ml cholera toxin (Sigma-Aldrich), 100 units/ml penicillin, and 100 μg/ml streptomycin (Sigma-Aldrich).

All cells were maintained at 37°C in 5% CO_2_ and were regularly tested for mycoplasma contamination.

### Drugs

Losmapimod, previously known as GSK-AHAB (*26*) (BioNova, 331-21264-1) was a p38α and β inhibitor. BJhTERT cells were treated at a concentration of 10 µM for an hour before the exposition to the 5 Gy challenging dose. Then, it was incubated for an additional hour before being fixed.

### Cell transfection

Cells were seeded into 10-cm plates overnight to reach 70% confluency before transfection. hsa-miRNA mimics, hsa-miRNA inhibitors and negative and positive controls were obtained from mirVana and siRNA duplexes were obtained from Dharmacon (Supplementary Table 2). Transfections were performed using RNAiMax Lipofectamine Reagent Mix (Life Technologies), according to the manufactureŕs instructions.

### Cell viability assay

Cells were seeded in 6-well plates at appropriate dilutions in triplicates and treated with ionizing radiation, as indicated in the corresponding figure legends, the following day. After 14-16 days, colonies were stained with a solution containing 0.5% crystal violet and 20% ethanol, followed by several washes with water. Crystal violet was then resolubilized with 10% acetic acid and absorbance (595 nm) was measured using the Varioskan Flash microplate reader (Thermo Electron corporation).

### Neutral comet assay

Following exposure to the indicated doses of ionizing radiation, cells were trypsinized, washed and resuspended in ice-cold PBS. Over 10^4^ cells were mixed with low-melting-point agarose (1%) at a ratio of 1:1 (v/v) by pipetting up and down and immediately pipetted onto an agarose-coated microscope slide. The slide was laid flat at 4°C for 30 minutes and then incubated overnight in lysis buffer (2.5 M NaCl, 100 mM EDTA, 10 mM Tris-HCl pH 7.5 and 200 mM NaOH). Next day, slides were immersed in neutral electrophoresis solution (100 mM Tris-HCl and 300 mM sodium acetate) for 30 minutes. Then, slides were placed in an electrophoresis unit containing electrophoresis solution and a voltage of 1 volt per cm was applied for 45 minutes at 4°C. Excess electrophoresis solution was drained, and slides were fixed in 70% ethanol for 30 minutes at room temperature, dried for 15 minutes at 37°C, and stained with SYBR Gold for 30 minutes at room temperature in the dark. Images were acquired using a Leica DMI8 Thunder fluorescence microscope and the analysis of comet tail length was measured using the OpenComet plug-in of the FIJI software.

### Immunofluorescence

All the experiments were performed in close collaboration with the microscopy unit at CABIMER.

Cells were seeded on coverslips. For RPA visualization, cells were treated with pre-extraction buffer (25 mM Tris-HCl, pH 7.5, 50 mM NaCl, 1 mM EDTA, 3 mM MgCl_2_, 300 mM sucrose and 0.2% Triton X-100) for 5 minutes on ice to pre-extract soluble proteins. Then, cells were washed once with PBS and fixed with 4% paraformaldehyde in PBS for 15 minutes. For γH2AX and phospho-P38 (Thr180-Tyr182) foci, cells were fixed with 4% paraformaldehyde in PBS for 15 minutes and permeabilized with 0,5% PBS-Triton for 15 minutes. For 53BP1 visualization, cells were fixed with ice-cold methanol for 10 minutes, followed by incubation with ice-cold acetone for 30 seconds.

Followed the fixation, cells were washed twice with PBS, and blocked for 1 hour with 5% FBS in PBS, co-stained with anti-RPA (Abcam #ab21752), anti-53BP1 (Novus Biologicals #NB100-304), anti-γH2AX (Abcam #ab22551) and anti-phospho-P38 (Thr180-Tyr182) (Protein Tech #28796-1-AP) primary antibodies diluted in blocking buffer overnight, washed again with PBS and then co-immunostained with the secondary antibodies anti-mouse Alexa Fluor 594 (Invitrogen #A11032) for RPA and γH2AX or anti-rabbit Alexa Fluor 488 (Invitrogen #A11034) for 53BP1 and phospho-P38 (Thr180-Tyr182) in blocking buffer for 1 hour at room temperature. Coverslips were then mounted into glass slides using Vectashield mounting medium with DAPI (Vector laboratories). Images were acquired using a Leica DMI8 Thunder fluorescence microscope. In all of them, 63x microscope objective was used for sample visualization and image acquisition. The analysis of the number of foci formation and the intensity was performed automatically using FIJI software.

### Cell cycle analysis

Cells were harvested by tripsinization, centrifugated at 500g for 5 minutes and washed once with PBS. Cells were then fixed with 1 ml of cold 70% ethanol (Merck) overnight at 4°C. Next day, ells were then centrifugated at 500 g for 5 minutes, washed with PBS and incubated with 250 µg/ml RNAse A (Sigma) and 10 µg/ml propidium iodide (Fluka) in PBS for 30 minutes at 37°C in the dark. Cells were analysed using a LSRFortessa X-20 Flow Cytometer (BD Biosciences). At least 10^4^ events were recorded for each sample. Cell cycle distributions were further analysed using ModFit LT 5.0 software (Verity Software House Inc).

### UV laser micro-irradiation

Cells were micro-irradiated using a wide-field Angström’s microscope (Leica) equipped with a Micropoint pulsed dye laser of 365 nm (Photonic Instruments, Inc.). Cells were seeded in 4-well plates. Next day, about 40–50 cells were micro-irradiated with one laser stripe per cell. For gH2AX study, cells were fixed with 4% paraformaldehyde in PBS for 15 minutes, permeabilized with 0,5% PBS-Triton for 15 minutes, and blocked for 30 minutes with 5% bovine serum albumin (BSA) in PBS. Anti-γH2AX primary antibody (Abcam #ab22551) was diluted in 1% BSA in PBST (PBS containing 0.01% Tween-20) and incubated for 1 hour, washed with PBS and then co-immunostained with the secondary antibody anti-mouse Alexa Fluor 594 (Invitrogen #A11032) diluted in 1% BSA in PBST for 1 hour at room temperature. After several washes, cells were incubated with DAPI to visualize the nucleus. Images were acquired using a Leica DMI8 Thunder fluorescence microscope. In all of them, 63x microscope objective was used for sample visualization and image acquisition. Quantitative analyses of the stripes were carried out in random areas using FIJI software. The number of stripes was quantified in 20–30 cells per preparation.

### Proximity ligation assay (PLA)

PLAs were performed using the Duolink PLA Kit (Duolink) according to the manufacturer’s protocol. Briefly, BJhTERT were treated with ionizing radiation, incubated 1 h and then collected. Coverslips were washed with PBS, fixed on ice with methanol for 10 min followed by acetone for 30 s, washed three times with PBS and blocked with blocking solution from the Duolink PLA Kit for 30 min at 37 °C. Samples were incubated with primary antibodies against CCAR2 (Cell Signalling #5857) and p38 (Cell Signalling #8680) overnight at 4°C, followed by MINUS and PLUS secondary PLA probes (anti-mouse minus and anti-rabbit plus) for 1 h at 37 °C. Detection was carried out with the Duolink Detection Kit Red (Olink Bioscience). At least 200 cells were analysed using a Leica DMI8 Thunder fluorescence microscope and foci counted automatically using using FIJI software.

### Immunoprecipitation

Cells were harvested in lysis buffer (50 mM Tris–HCl, pH 7.4, 150mM NaCl, 1mM EDTA, 0.2 % Triton X-100, 1X protease inhibitors (Roche), 1X phosphatase inhibitor cocktail 1 (Sigma)) and incubated for 30 min on ice with Benzonase (90 U/ml). Protein extract (2 mg) was incubated at 4°C with 10 μl of anti-CCAR2 or anti-P38 antibodies or with an equivalent amount of IgG (Mouse or Rabbit) as the negative control. Afterward, extracts were incubated with magnetic protein A (CCAR2) or G (P38) Dynabeads (Novex) overnight. Beads were then washed three times with lysis buffer, and the precipitate was eluted in a mixture of 10 μl of Laemmli buffer 4x and 30 μl of lysis buffer.

### Extracellular vesicles isolation and characterization

For sEVs isolation, 10^5^ cells were seeded into three 15-cm plates per condition. After 3-5 days, culture media was centrifuged at 500 g for 10 minutes to remove any cell contamination and filtered through a 1.2 µm Whatman puradisc 13 syringe filters (WAH68221312). Following concentration of the supernatant with an Amicon Ultra 15 ml centrifugal filter (UFC9100), extracellular vesicles (sEVs) were isolated using the exoRNeasy Midi Kit (Qiagen #77144) following the manufacturer’s instructions.

For sEVs tracking, lipophilic membranes were stained by adding 1 µM Dil (1,1’-Dioctadecyl-3,3,3’,3’-Tetramethylindocarbocyanine Perchlorate, Invitrogen) to the culture media of both non-pre-irradiated and pre-irradiated cells and incubated for 1 hour at 37°C. Next, the media from non-pre-irradiated and pre-irradiated cells were exchanged with Dil-labelled media from either pre-irradiated or non-pre-irradiated cells, and the cultures were incubated for an additional 3 hours. Cells were then fixed with 4% paraformaldehyde (w/v) in PBS for 15 minutes. Followed fixation, cells were washed twice with PBS and coverslips were then mounted into microscope slides using Vectashield mounting medium with DAPI (Vector laboratories). Images were acquired using a Leica DMI8 Thunder fluorescence microscope using the 63x objective.

For electron microscopy visualization, purified sEVs were adhered to uncoated copper grids and stained with 4% uranyl acetate for 4 minutes. Images were acquired using a ZEISS LIBRA 120 transmission electron microscopy. Diameter, roundness, perimeter and area of sEVs were measured using TEM ExosomeAnalyzer software (*32*) For flow cytometric analysis, purified sEVs were analysed by using the ExoStep CD81/CD9 flow detection reagent (Immunostep ExoS-25-G81; ExoS-25-G9) following the manufacturer’s instructions. Samples were analysed using a LSRFortessa X-20 Flow Cytometer (BD Biosciences). At least 10^4^ CD63 positive events were recorded for each sample.

### sEVs RNA extraction and qRT-PCR

miRNA was extracted using RNeasy Micro Kit (Qiagen #74004) or miRNeasy Micro Kit (Qiagen #217084) according to the manufacturer’s instructions, followed by the digestion of genomic DNA by DNase treatment. To generate cDNA from the extracted RNA, miRCURY LNA RT Kit (Qiagen #339340) was used following the manufacturer’s protocol.

The cDNA template was used to quantify miRNA expression employing the miRCURY miRNA PCR assay for hsa-miR-126-3p, hsa-miR451a, hsa-miR-1-3p (positive control) and hsa-miR-103-3p (housekeeping) (Qiagen #YP00204227, #YP02119305, #YP00204344, #YP00204063). The reaction mix setup was performed according to manufacturer’s recommendations. The PCR cycling conditions consisted of an initial heat activating step at 95°C for 2 minutes, followed by 45 cycles of denaturation at 95°C for 10 seconds, annealing at 56°C for 60 seconds and extension at 56°C for 60 seconds.

### sEVs RNAseq analysis

All sequencing was performed at the Genomics Unit of CABIMER.

miRNA purification was performed as explained above. The quality and integrity for each RNA sample were verified using a Bioanalyzer 2100 instrument (Agilent) before proceeding to the RNA-Seq protocol. Small RNA libraries were constructed with the Small RNA Sequencing Kit v3 for Illumina Platforms (NEXTflex) according to the manufacturer’s protocol. Libraries were then sequenced on a NovaSeq6000 (Illumina) to generate 100-base single-ends reads.

For RNA-seq processing, Galaxy web platform was used (usegalaxy.org) (*33*) First, single-end reads were quality controlled by removing adapter sequences, low-quality bases and short reads using FastQC and Trimmomatic tool. Next, HISAT2 program was employed to align cleaned RNA-seq paired end reads and finally, the gene expression measurement of the aligned sequences was done by FeatureCounts tool. Data was analysed with Limma tool, considering p-value<0.05 and fold change <-2 and >2 (*34*). Genes were considered as differentially expressed based on the comparison of the relative abundance between both groups (pre-irradiated sEVs vs non-pre-irradiated sEVs).

GO enrichment analysis was performed using ShinyGO 0.81.

### sEVs Mass Spectrometry sample preparation

Purified sEVs were resuspended in RIPA lysis buffer (Thermo Scientific) by gently pipetting and vortexing. Then, lysates were incubated with 10 mM TCEP (Merck) for 45 min at 37°C with shaking to promote disulfide reduction, followed by the cysteine alkylation with 40 nM cloracetamide (Merck) for 30 min at room temperature in the dark. Meanwhile, beads were prepared using paramagnetic bead technology based on the Single-Pot Solid-Phase-Enhanced Sample Preparation (SP3), in which two types of SpeedBeads, mixed at a 1:1 ratio, were used: 45152105050250 and 65152105050250 (Cytiva). Samples were incubated with the beads mixture and 100% acetonitrile (Merck) with shaking for 35 min at room temperature. Proteins were then washed twice with 70% ethanol (Merck), followed by two additional washes with 100% acetonitrile. Finally, the beads were incubated with 250 ng trypsin in 50 mM ammonium bicarbonate at 37°C with shaking overnight.

Digested peptides were separated from the beads using a 0.45µm centrifugal filter unit (Millipore) and acidified by adding 2% trifluoroacetic (TFA) acid. Subsequently, peptides were desalted and concentrated on triple-disc C18 Stage-tips as previously described (*39*). Stage-tips were in-house assembled using 200 µL micro pipet tips and C18 matrix (Sigma-Aldrich). Stage-tips were first activated by passing through 100[µl of methanol. Next, 100[µl of Buffer B (80% acetonitrile, 0.1% formic acid), 100[µl of Buffer A (0.1% formic acid), the acidified peptide sample, and two times 100[µl

Buffer A were passed through the Stage-tip. Elution was performed twice with 30 µl of Elution buffer (32,5% acetonitrile, 0.1% formic acid solution). Samples were vacuum dried using a Universal Vacuum System UVS400S coupled to a SpeedVac SPD121P (Thermo) and stored at −20[°C. Prior to mass spectrometry analysis, samples were reconstituted in 20 µl 0.1% formic acid and transferred to autoload vials.

### Mass Spectrometry Data acquisition

Mass spectrometry-based proteomics data was acquired at the Proteomics facility in Institute of Biomedicine of Seville (IBiS). As previously done (*40*), samples were measured using a nanoElute II LC system coupled to a timsTOF SCP mass spectrometer with an electrospray source (Bruker Daltonics). LC separations were performed on C18 HPLC column (Aurora 25 cm and 75 µm ID, IonOpticks) kept at 50°C. Gradient elution was performed with a binary system consisting of (A) 0.1% aqueous formic acid and (B) 0.1% formic acid in Acetonitrile. An increasing linear gradient (v/v) was used (t (min), %B): (0, 2); (40, 17); (60, 25); (66, 37); (67, 95), followed by an equilibration step.

Mass spectrometric analysis was performed in a data independent acquisition parallel accumulation serial fragmentation (dia-PASEF) mode, with 100–1700 m/z mass range, an ion mobility range from 0.64 to 1.45[V[s[cm−2, capillary voltage set to 1500 V, an accumulation and ramp time at 100 ms and the collision energy as a linear ramp from 20[eV at 1/K0[=[0.6[V[s[cm−2 to 59[eV at 1/K0[=[1.6[V[s[cm^−2^.

### Mass Spectrometry Data analysis

Mass spectrometry raw dia-PASEF data was analyzed using Biognosys Spectronaut (v19.5.241126) using a directDIA+ search with modified BGS Factory settings. Predicted spectral library was built using a FASTA file corresponding to the reference human proteome (Uniprot 19^th^ January 2024) only including canonical proteins (20,413 entries). Maximum number of variable modification were set to 2 and Carbamidomethyl (C) was disabled as a fixed modification. A pivot report was generated and further processed in the Perseus Computational Platform (*41*). Proteins not identified in every replicate for at least one condition were removed and missing values were randomly imputed using normally distributed values with 0.3 width and 1.8 down shift considering the total matrix. Statistical analysis was performed using two-sided Student’s t tests and exported to MS Excel for comprehensive browsing of the data.

### Statistical analysis

Statistical significance was determined with the test indicated in the corresponding figure legend using PRISM software (Graphpad Software). Statistically significant differences were labelled with one, two, three or four asterisks if p<0.05, p<0.01, p<0.001 or p<0.0001, respectively. Specific replicate numbers (n) for each experiment can be found in the corresponding figure legends.

