## Supplementary figures and images for "miRNA-mediated cell-to-cell communications boost DNA repair during the Radioadaptative Response"

### Supplemental Figure 1

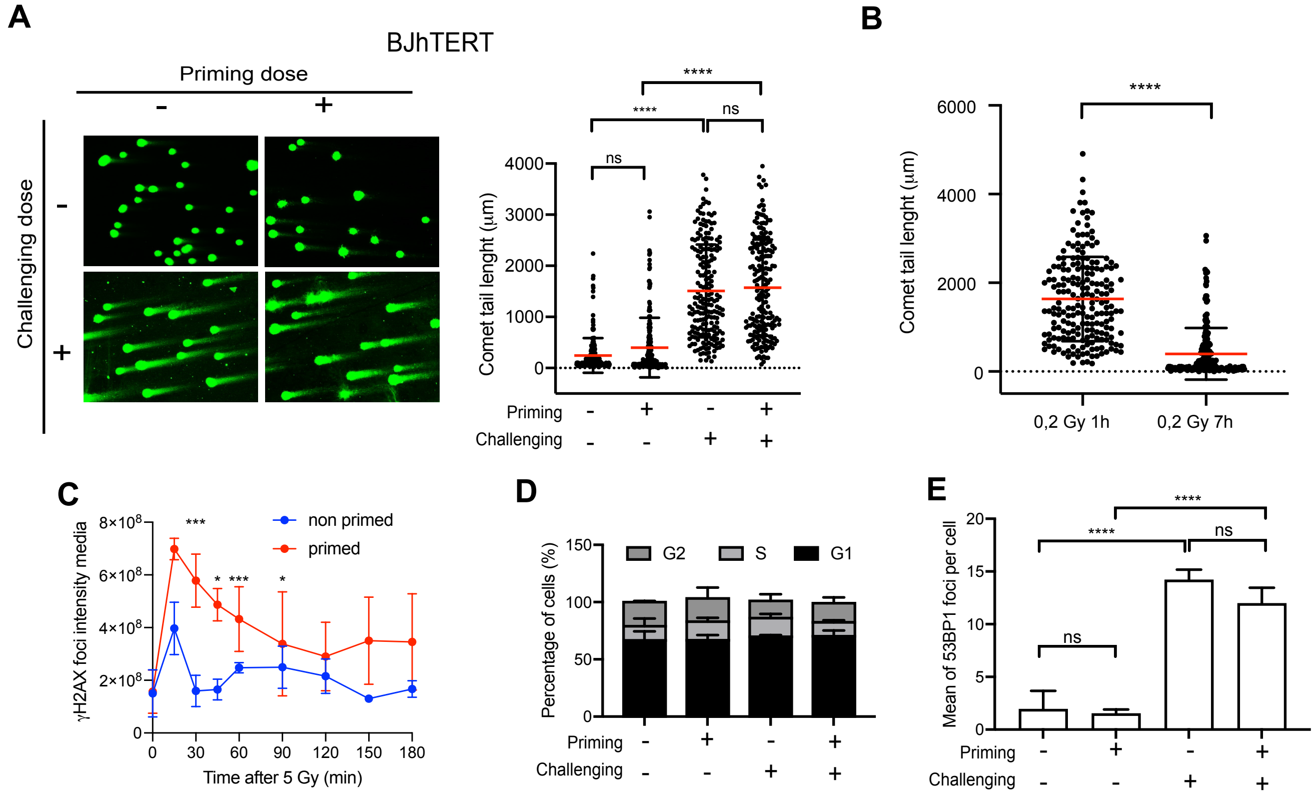


Dominguez-Perez Supplementary Figure 1

### Supplemental Figure 2

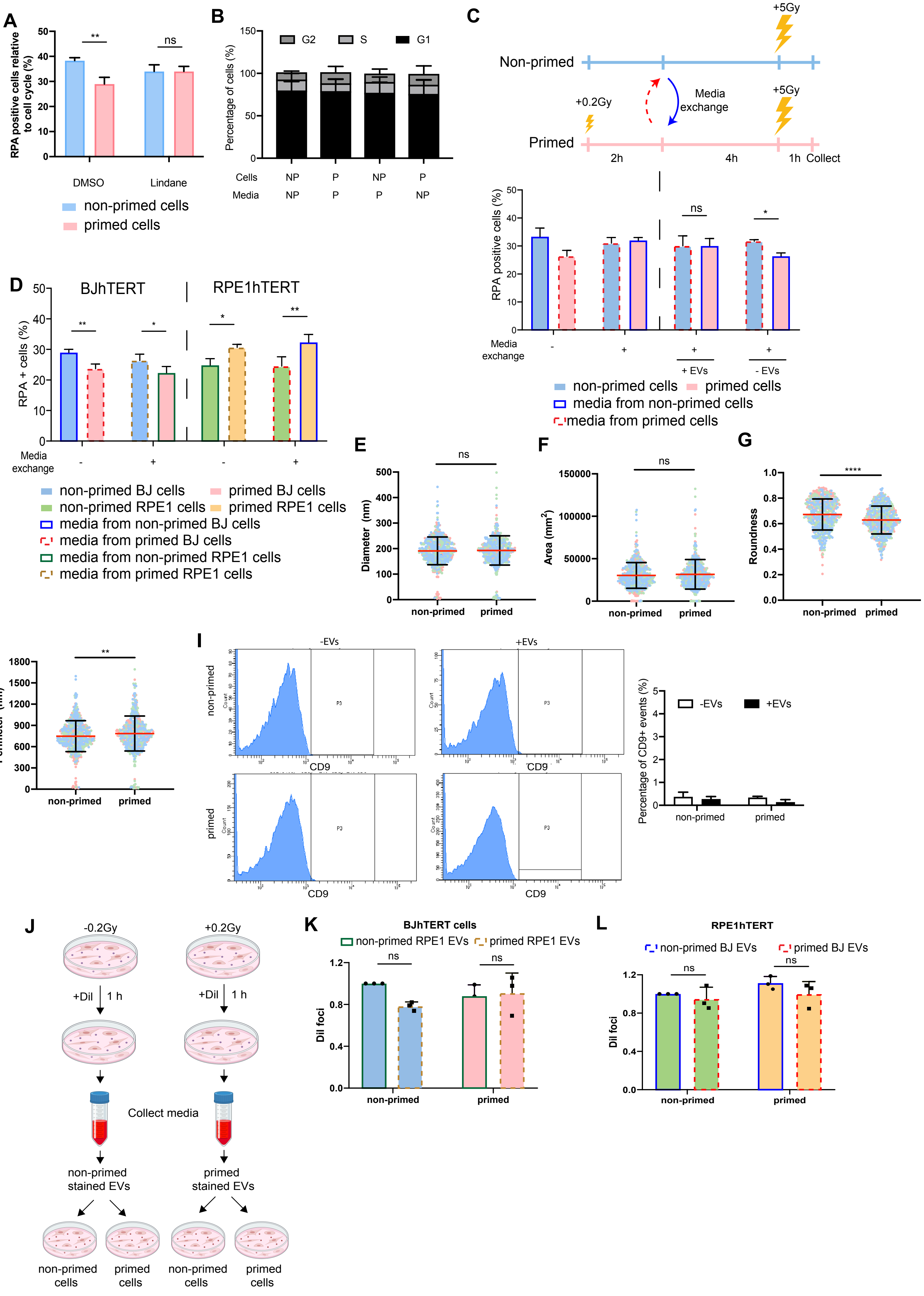

### Supplemental Figure 3

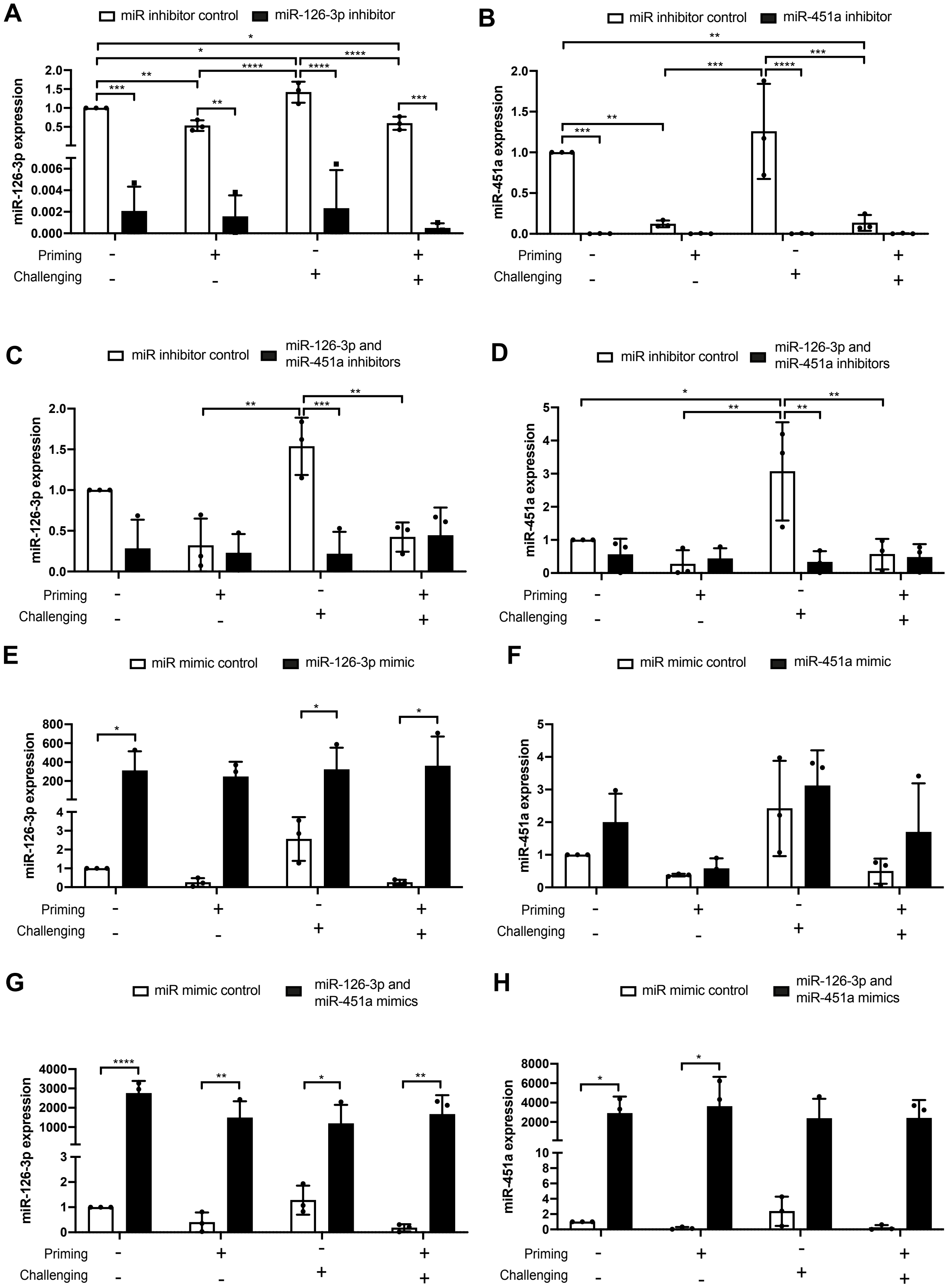


Dominguez-Perez Supplementary Figure 3

### Supplemental Figure 4

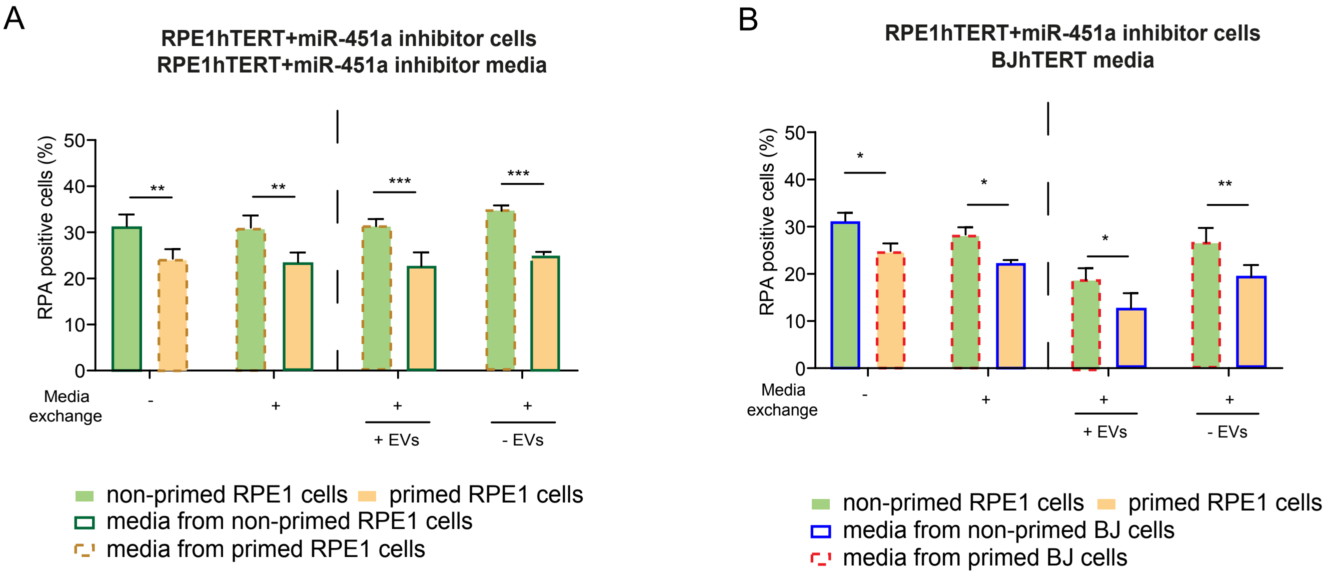

### Supplemental Figure 5

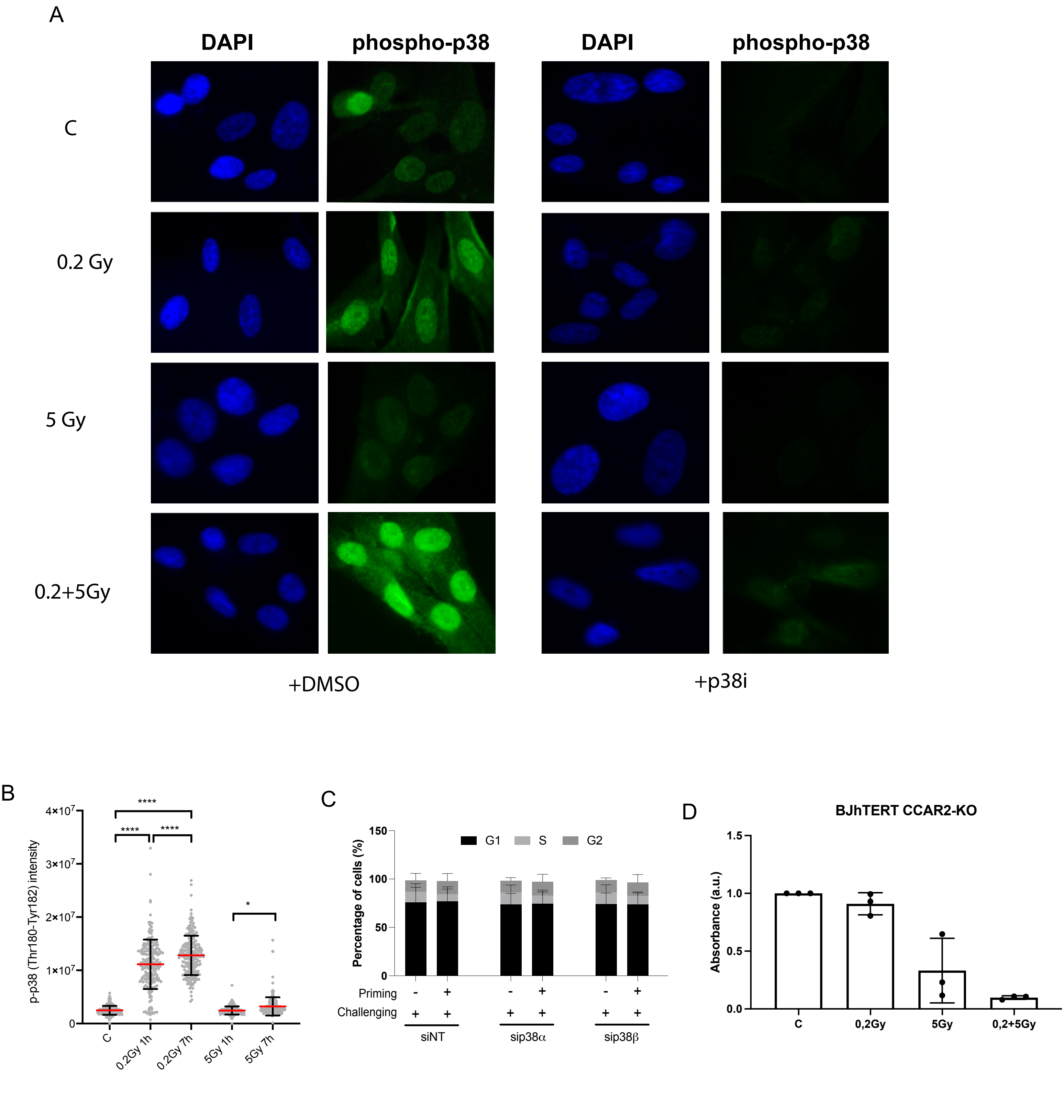


Dominguez-Perez Supplementary Figure 5
